# Epigenome analysis of an algae-infecting giant virus reveals a unique methylation motif catalogue

**DOI:** 10.1101/2025.08.10.669558

**Authors:** Alexander R. Truchon, Erik R. Zinser, Steven W. Wilhelm

**Author notes:** Corresponding author: Steven W. Wilhelm.

## Abstract

DNA methylation can epigenetically alter gene expression and serve as a mechanism for genomic stabilization. Advancements in long-read sequencing technology have allowed for increased exploration into the methylation profiles of various organisms, including viruses. Studies into the *Nucleocytoviricota* phylum of giant dsDNA viruses have revealed unique strategies for genomic methylation. However, given the diversity across this phylum, further inquiries into specific lineages are necessary. *Kratosvirus quantuckense* is predicted to encode six distinct methyltransferases, which bear homology to other methyltransferases across the many clades of *Nucleocytoviricota*. We found that this virus methylates its own DNA with high consistency and targets up to nine different motifs for DNA adenine methylation. Methylation levels varied depending on the associated motif. Likewise, distinct motifs were enriched within unique genomic regions. Collectively this suggests that each methyltransferase targets unique DNA regions and may suggest they have varying functionality. This work reveals an array of methyltransferase activity in *Kratosvirus quantuckense* and begins to implicate the importance of DNA methylation to the *Nucleocytoviricota* infection cycle.

## INTRODUCTION

The role of DNA methylation has been characterized largely through robust genetic analysis in a handful of model organisms (1–4). DNA methyltransferases (MTases) catalyze the addition of a methyl group typically onto either an adenine, yielding N^6^-methyladenine (6mA), or a cytosine, yielding often 5-methylcytosine (5mC) in either a CpG or non-CpG context, and less often N^4^-methylcytosine (4mC) or 5-hydroxymethylcytosine (5hmC) (4–7). These modifications can have significant effects on the functional potential and structure of the organism’s genome. While 5mC methylation has been classically attributed to repressing gene promoter activity, particularly in eukaryotes (8), 6mA methylation has been shown to exhibit a myriad of functions in prokaryotes (9). In addition to regulating gene expression (10), DNA adenine methylation (6mA) has been largely associated with restriction modification (RM) systems that in tandem methylate an organism’s DNA while targeting unknown DNA for degradation *via* restriction endonuclease activity (5, 9, 11, 12). While RM systems have not been identified in eukaryotes, evidence of 6mA methylation has been identified in single-celled eukaryotes (13, 14), which may point to acquisition of methylation genes *via* horizontal gene transfer or even activity of symbiotic bacteria (15). 6mA methylation has also been associated with DNA replication and mismatch repair (12).

Few analyses into the *Nucleocytoviricota* have delved into the methylation of viral genomes, despite a high abundance of DNA MTases encoded within their genomes (16, 17). Of particular significance is Paramecium bursaria Chlorella Virus 1 (PBCV-1), which encodes five predicted DNA MTases, two of which have strongly defined functionality (16, 18, 19). These two adenine-specific MTases, M. *Cvi*AI and M. *Cvi*AII, are flanked by restriction endonucleases which target GATC and CATG cut sites, respectively (18). Methylation of the virus’s own genome provides protection from these self-encoded restriction enzymes while allowing for degradation of host DNA, similarly to the Dam RM systems typical of bacteria which target invading phage DNA. For this reason, a high proportion of GATC or CATG sites are fully methylated on each strand, with most of the remaining sites being hemi-methylated (16). Given the importance of evading restriction endonucleases for proper replication of the viral genome, universal genomic methylation is an expected functionality of these types of MTases.

The three other DNA MTases encoded by PBCV-1 have not yet been shown to be functional, though if they are, they likely target cytosine sites rather than adenine (18). They are also not genomically colocalized with known restriction enzymes. This is the case for many DNA MTases encoded by the *Nucleocytoviricota* (17), thus bringing their function into question. Several DNA MTases were characterized in the Pandoraviruses, a lineage of *Nucleocytoviricota* with particularly large genomes (20), which methylate a variety of cytosine motifs and bear high phylogenetic identity to *Acanthamoeba spp.* MTases, suggesting these are genes are host derived (17). However, *Mollivirus sibericum, Cedratvirus kamchatka,* and several Marseilleviruses encode for adenine-specific MTases and do methylate their own genomes (17). In Marseilleviruses in particular, this appears to again be associated with an RM system (17, 21). Outside of this information the role of DNA methylation in viral infection is still largely unclear.

This is particularly significant regarding *Kratosvirus quantuckense*, a virus of the *Nucleocytoviricota* which infects the eukaryotic brown alga *Aureococcus anophagefferens* (22, 23). *K. quantuckense* encodes for several DNA-specific MTases (22). However, these MTases are not homologous to the functionally defined MTases of PBCV-1, nor do they co-occur with any identifiable restriction endonucleases. The high density of MTases encoded by *K. quantuckense* merits further analysis of their function during infection, their activity on genomic viral DNA, and the variability of sites that are targeted for methylation. Moreover, the diversity among these MTases may imply divergent functionality, possibly involving genomic stability within the virocell and regulation of DNA packaging.

Here we used Nanopore long-read sequencing to define the methylation landscape of *Kratosvirus quantuckense* strain AaV. Repeated whole genome sequencing of four biological replicates of viral DNA revealed consistent methylation patterns of both adenines and cytosines. Analyses of these methylation patterns across the genome revealed the presence of several motifs which are likely targeted by different MTases encoded by AaV. We present this information in the context of virus-host interactions, the diversity of methylation strategies across the *Nucleocytoviricota*, and how this generally overlooked aspect of the viral genome shapes the potential success of this pathogen.

## METHODS

### Culture Conditions

Three 750 mL batch cultures of non-axenic *Aureococcus anophagefferens* CCMP1984 were grown in ASP_12_A media (24) on a 12:12 light:dark cycle at 19° C and an irradiance of ∼90 µmol photons m^-2^ s^-1^. After one week of logarithmic growth, cultures were diluted 10:1 in ASP_12_A to 1 L to prevent nutrient starvation. Two days following dilution, cultures were infected with 10 mL of fresh *K. quantuckense* strain AaV in ASP_12_A (22). Fresh AaV had been prepared through infection of a 25 mL culture of *A. anophagefferens* CCMP1984 with AaV, after which lysate was pushed through a Durapore 0.45-µM nominal pore-size PVDF-membrane syringe filter (MilliporeSigma; Burlington, MA) before inoculation. Once infected cultures were completely lysed (∼3 d), lysate was stored at 4° C.

### Preparation of Viral Particles for DNA Extraction

Lysate was prefiltered through an Isopore™ 0.4-µM nominal pore-size 25-mm diameter polycarbonate membrane syringe filters (MilliporeSigma; Burlington, MA) to remove lysed cellular material and surplus heterotrophic bacteria. Each liter of filtered lysate was then concentrated by tangential flow filtration (TFF) as previously described (25). Briefly, lysate was sequentially concentrated with a Pellicon XL 30 kDa Cassette (MilliporeSigma; Burlington, MA) using a Labscale TFF System (MilliporeSigma; Burlington, MA). Lysate was concentrated from 1 L to approximately 25 mL, yielding a theoretical 40-fold increase in viral particle concentration. Concentrated lysate was once again filtered through a Durapore 0.45 µm pore-size PVDF membrane syringe filter (MilliporeSigma; Burlington, MA). Lysate was enumerated on a CytoFLEX flow cytometer (C07821) by gating on the Violet Side Scatter channel using a 405 nm violet laser (26) to ensure production of viral particles. To further clean AaV particles, lysate was centrifuged at 2,000 xG for 5 min to pellet remaining bacterial cells. The supernatant was moved to a clean tube and 10% Triton X-100 was added at a final concentration of 1%. To concentrate particles, the supernatant was then centrifuged at 29,000 rpm for 75 min to pellet viral particles, which were subsequently resuspended in 400 µL ASP_12_A.

### DNA Extraction and Sequencing

Lysozyme (120 µL of 20 mg/mL) and 1 µL RNase (10 mg/mL) were added to the concentrated viral particles and incubated at 37° C for 30 min. Viral particles were then treated with 10 µL of proteinase K solution (20 mg/mL, 3 mM CaCl_2_, 200 mM Tris buffer) along with 40 µL lysis solution (0.5% SDS, 10 mM EDTA, 20 mM sodium acetate) and incubated at 55°C for 2 hours while gently shaking. DNA was extracted using a standard phenol-chloroform method. DNA concentration and purity was determined on a NanoDrop spectrophotometer and size of extracted DNA was visualized using gel electrophoresis.

DNA extracts from each replicate were individually sequenced using a new MinION Mk1B sequencing device fitted with a Flongle adapter for Flongle Flow Cells (R10.4.1) (Oxford Nanopore Technologies; Oxford, UK). Sequencing libraries were prepared according to manufacturer’s instructions using a V14 Ligation Sequencing Kit (SQK-LSK114) and the Flongle Sequencing Expansion (EXP-FSE002) (Oxford Nanopore Technologie; Oxford, UK).

To serve as a control, DNA was extracted from a fourth biological replicate of concentrated viral lysate. This DNA was diluted to approximately 3 ng/µL before performing whole genome amplification (WGA) with the NEB phi29-XT WGA Kit (New England Biolabs, Ipswich, MA). This WGA DNA and an aliquot of the original, undiluted DNA from the fourth replicate were then sequenced on tandem Flongle Flow Cells (R10.4.1). All sequencing runs performed for this analysis are summarized in **Table S1**.

### Methylation Calling and Initial Analysis

POD5 files generated from long-read sequencing were aligned to the most recent version of the AaV genome (27) and called for 5mC and 6mA methylation using Bonito basecaller with the dna_r10.4.1_e8.2_400bps_hac@v5.0.0 model with a minimum q-score of 9 (28). Called and aligned reads were sorted and indexed using Samtools (29). BedMethyl tables, which display total methylation at all genomic sites, were created using the Modkit pileup command (Oxford Nanopore Technologies). To avoid considering methylation at sites with only a small number of reads mapped (*i.e.*, single nucleotide polymorphisms) python scripts were used to filter the bedMethyl file down to only sites that contain the respective nucleotide (either adenine for 6mA methylation or cytosine for 5mC methylation). Methylation frequency (*i.e.* proportion of nucleotides at a given site methylated) for each individual site was calculated using the equation:

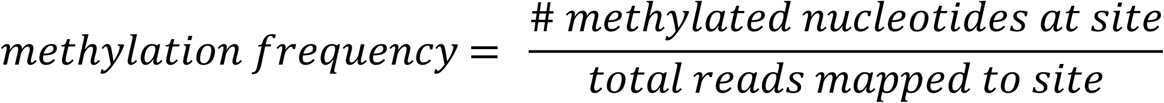

Likewise, genomic methylation fraction was calculated by determining the proportion of all sites in the genome that meet the threshold to be considered methylated, which, for the purpose of this study, is 70% methylation frequency. For methylated site enrichment scores of specific genomic regions, every 1000 bp frequently methylated sites were counted within a sliding window of 5000 bp before being normalized by the total number of the respective site within the region. Z-scores from enrichment scores were calculated and used for Circos heatmaps.

To determine WGA-corrected methylation frequency for each site, methylation frequency scores from the WGA-amplified library BedMethyl files were subtracted from the methylation frequency scores from the respective unamplified library. Python scripts were used to determine the frequency of nucleotides surrounding highly methylated sites (>80% for 6mA and >50% for 5mC). Methylation maps were generated using python scripts and imaged using Circos.

### Methylation Targeted Motif Identification and Characterization

To identify DNA sequences likely to be targeted for methylation in the AaV genome, corrected BedMethyl tables were run through the motif detection software Nanomotif using the motif discovery function. This process was performed for all three libraries and any motifs identified were retained for future analysis. Overall, nine putative targeted motifs were discovered, five of which were palindromic. For palindromic motifs, python scripts were used to pair scores belonging to the same palindrome on opposite strands to determine hemi-methylation of these sites.

To identify genomic regions in which specific motifs are enriched, the web server DistAMo was used to visualize motif distribution. Coding regions that were overrepresented with a specific motif with a Z-score >= 2 were identified for each motif. Genes that were found to be associated with a specific motif were clustered using Cytoscape where every line connects a gene to any motifs that are overrepresented in the region.

### Phylogenetic Analysis of Viral Methyltransferases

To better characterize the DNA MTases, genes were called from sequenced *Nucleocytoviricota* genomes (including AaV) using the Nanomotif tool MTase-linker, which compares coding regions of each respective genome to the entire REBASE database and flags likely DNA MTases. The number of DNA MTases identified through this process was normalized to both the length of the respective virus’s genome as well as the number of identified coding regions for comparison across all viruses.

To generate a phylogenetic framework of *Nucleocytoviricota* DNA MTases, protein sequences for each gene were aligned using MAFFT v7.520 using a maximum number of iterative refinements of 1000. Sequence alignments were then trimmed using trimAl v1.4.rev15 with a gap threshold of 0.9 and a 25% conservation. A maximum-likelihood protein tree was constructed using IQ-TREE version 2.2.0.3 with 1000 bootstraps and visualized in IToL.

### Restriction Enzyme Digestion

To verify results noted from motif analysis, three restriction enzymes were used to verify the presence of methylation on AaV adenines. These included Hpy166II, which targets the GTNNAC motif, XhoI, which targets the CTCGAG motif (*i.e.* a proxy for CTNNAG), and XbaI, which targets the TCTAGA motif (*i.e.,* a proxy for CTAG). DpnI and DpnII were used as controls, which are expected to digest methylated and unmethylated motifs respectively. All digestions were performed with either AaV, whole genome amplified AaV, or *A. anophagefferens* DNA in rCutSmart Buffer (New England Biolabs, Ipswich, MA) for 15 min at 37° C. Reactions were inactivated at 65° C for 20 min and then visualized *via* gel electrophoresis.

## RESULTS

AaV encode seven nucleotide-associated MTases, one of which is likely to act as an RNA methylase based on homologous sequences. Among the six remaining MTases, five appear in the Restriction Enzyme Database (REBASE), a catalog identifying restriction endonucleases and MTases (30). All are considered type II MTases, meaning they exist as distinct ORFs with no associated endonuclease domains. All have the characteristic domains of an MTase [*i.e.*, fgg, dppy, and the DNA target recognition domain (TRD)] (**Table S2**) (31, 32). Four of these MTases fit into the γ subclass, signifying a motif order of fgg-dppy-TRD, while the final MTase belongs to the β subclass, signifying a motif order of dppy-TRD-fgg (33). When characterizing the function of these MTases using the Nanomotif MTase-caller tool (34), the five genes described above are characterized as adenine-specific MTases, while an additional MTase was identified as potentially cytosine-specific. This final gene, which was not described in REBASE, contains three of the six characteristic domains of a cytosine-specific MTase (fgg, pc, and env), though it appears truncated and lacks the three terminal domains that are consistent with cytosine specific MTases (qrr, ix, and x) (35). This gene bears a high identity to two MTases encoded by the *A. anophagefferens* mitochondrion (59.48% amino acid identity), one of which is also truncated while the other contains all six motifs. As it is unclear if the protein product for this gene is functional, it will not be further analyzed in this study. Furthermore, none of the MTases appear to be packaged in the viral capsid (36), meaning they are likely only active after transcription initiation.

When compared to other giant viruses, AaV encodes more MTases in the context of the entire genome. Despite being one of the smaller *Nucleocytoviricota* with a genome size of just ∼380 kb, AaV encodes six MTases out of 384 ORFs, giving it a high ratio of MTases per genome (**Figure 1A**) and MTases per total encoded genes (**Figure 1B**) as compared to other similar viruses (**Table S3**). Of the viruses examined, these represent some of the highest rates of encoded DNA MTases, in a similar fashion to Ostreococcus lucimarinus Virus (OlV; 4 DNA MTases in a 190 kb genome), PBCV-1 (5 DNA MTases in a 330 kb genome), and Phaeocystis globosa Virus (5 DNA MTases in a 460 kb genome). While the larger *Nucleocytoviricota* like the pandoraviruses or Bodo saltans Virus do encode DNA MTases, none display the density seen in AaV (**Table S3**). Moreover, many amoebal *Mimiviridae*, like Acanthamoeba castellanii Mimivirus (APMV), do not appear to encode any traditional DNA MTases according to REBASE standards (**Table S3**).

**Figure 1.**
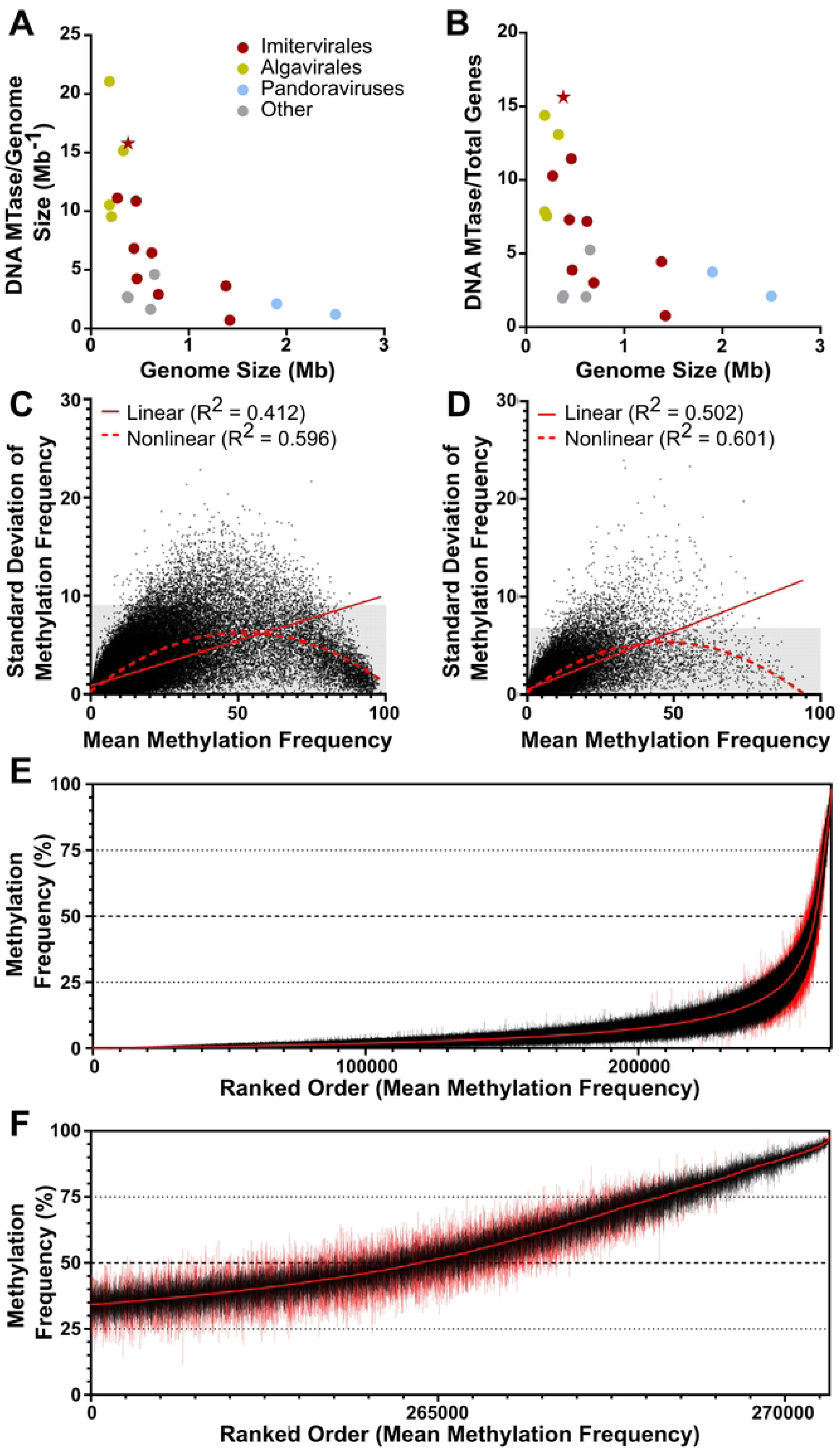
Base methyltransferase abundance and methylation occurrence in AaV. Encoded DNA methyltransferases of various *Nucleocytoviricota* normalized to genome size (A) and total coding potential (B). AaV is indicated with a star. Distribution of methylation frequency average scores and standard deviations (n=3) among every AaV adenine (C) and cytosine (D) within the AaV genome. Gray regions of the graph represent the lower 99% of standard deviations. R-squared statistics of linear and non-linear quadratic regressions are denoted. Adenines in AaV sorted by ranked order of lowest to highest mean methylation frequency (E-F). Mean methylation scores are denoted as the red line in the center, while lines extending from the center represent the range for each respective site. Sites within the 99^th^ percentile of standard deviation have red ranges. Figure 1F displays a magnified view of the most highly methylated sites in 1G.

Methylation frequency of both adenines (6mA) and cytosines in the AaV genome was consistent between each successive sequencing run (**Figure S1**). The standard deviation of the methylation frequency is lower than 10% for 99% of AaV adenines (**Figure 1C**) and lower than 7% for 99% of cytosines (**Figure 1D**). The standard deviation of the methylation frequency also follows a nonlinear quadratic regression in relation to the mean methylation frequency for both adenines and cytosines, showing that both low methylation and high methylation sites have reduced variability (**Figure 1C-D**). A vast majority of adenines have low methylation scores, *i.e.* below 25%, while the range of methylation scores for a given site increases as the mean increases (**Figure 1E**). However, while it is expected for there to be a higher standard deviation around larger scores, the absolute highest methylation frequencies have very consistent scores across all libraries (**Figure 1F**).

Genomic maps of the methylation of AaV are detailed in Figure 2. 6mA methylated adenines (**Figure S2A**) occur consistently throughout much of the genome at relatively even rates while 5mC methylated cytosines (**Figure S2B**) are frequently in distinct clusters. Several specific genes are highly methylated on adenine residues within the intragenic region, including the major capsid protein (MCP) in which 5.7% of adenines are methylated at a frequency over 75% (compared to 0.90% genome-wide) (**Figure 2A**). Highly methylated cytosines are frequently found in repeat containing proteins, one of which 9.1% of cytosines methylated at a frequency over 50% (compared to 0.63% genome-wide) (**Figure 2B**). Regardless, methylation enrichment occurs for both 6mA and 5mC methylation in different locations depending on the strand (**Figure S3**). On average, cytosines in the AaV genome are methylated at a rate of approximately 4.59% with 0.664% of cytosines being methylated at a rate higher than 50%. Adenines in the AaV genome are methylated at a rate of approximately 6.55% with 1.917% of adenines being methylated at a rate higher than 50%.

**Figure 2.**
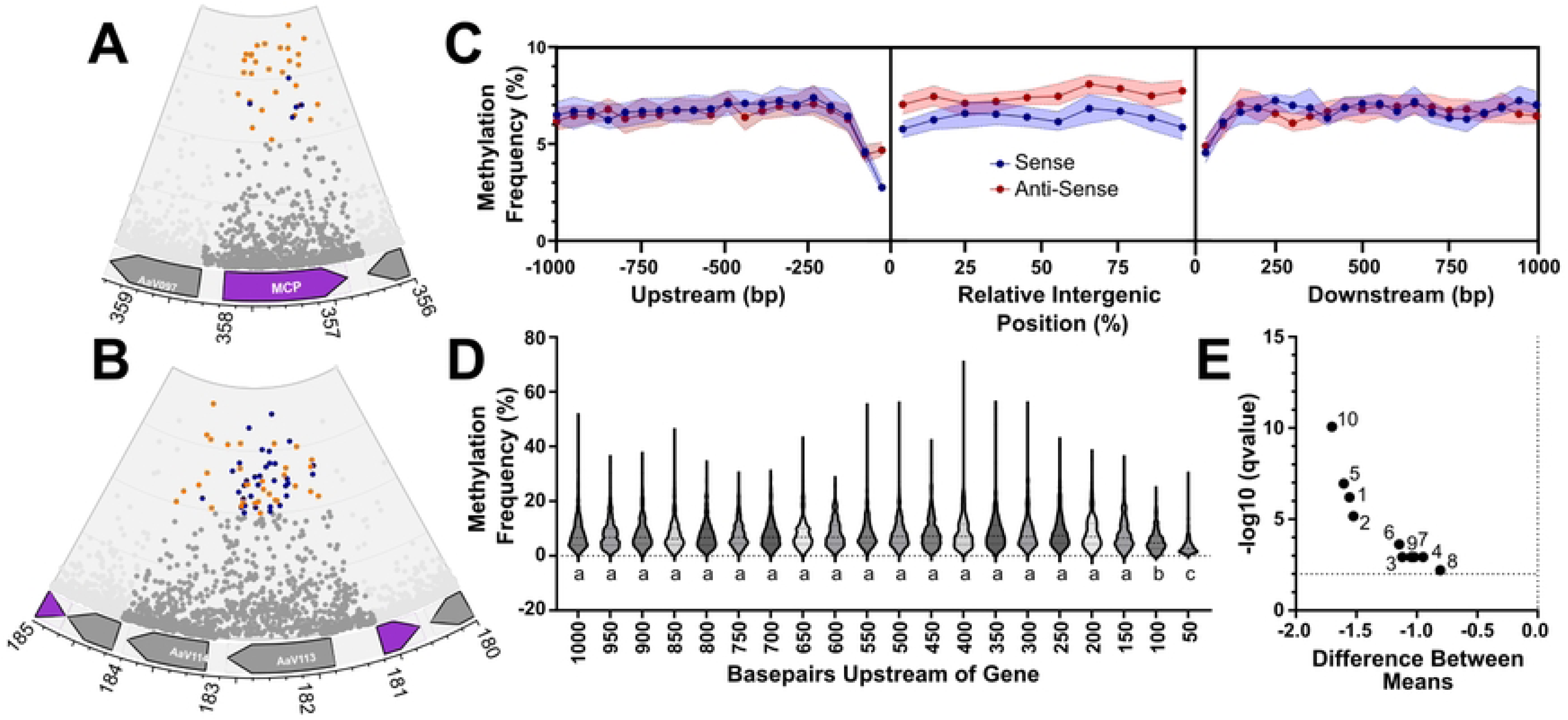
Trends in methylation frequency relative to coding regions. The major capsid protein shows an abundance of highly methylated adenines (A), while other regions are densely populated with methylated cytosines (B). The inner ring represents 100% methylation, and the outer ring represents 0%. Sites below 50% methylation frequency are shaded gray, cytosines above 50% are colored blue and adenines above 50% are colored orange. In 2A and 2B sites within coding regions outside those of interest are not shown. Genes on the sense strand are shown in purple while genes on the antisense strand are gray. Median methylation frequency is associated with intragenic regions along with the 1000 bp upstream and downstream regions for all coding regions of AaV (C), with 95% confidence intervals around the site shown in the respective color of the strand. Distribution of methylation frequencies in the upstream sense region for each gene is displayed in violin plots (D), with different letters above each plot signifying significant statistical difference (p<0.0001). The difference between averages of each normalized intragenic region is displayed in a volcano plot (F), with numbers representing the position along the gene (i.e., the terminal 5’ end represented by 1 and the terminal 3’ end represented by 10).

Methylation across intragenic regions, as well as the 1 kb upstream region and 1 kb downstream region for each gene were characterized collectively across the entire AaV genome. Intragenic regions for each gene were normalized into ten distinct regions, while each 50 bp window within the upstream and downstream regions were considered for methylation frequency, as per previous studies (37). Average methylation frequency for these sites was determined based on the total adenines a given region, and the averaged methylation scores across said region. Notably, methylation frequency across the gene body was consistently lower on the sense strand as compared to the anti-sense strand, which averaged 6-7% and 7-9% respectively (**Figure 2C,E**). Despite this, methylation did not vary largely within different intragenic regions (**Figure 2C**).

In regions both 100 bp upstream and 50 bp downstream of coding regions, a decrease in methylation frequency was visible (**Figure 2C**). This decrease was most notably exacerbated within the 50-bp immediately upstream of the gene body on the sense strand. This drop in methylation was significantly lower than the 19 other sites upstream of the gene body (**Figure 2D**). A similar pattern is notable on the antisense strand upstream of the gene body as well as on both strands downstream of the gene body, if not to a lesser extent.

The drop in methylation frequency immediately upstream of coding regions may be attributable to a promoter motif that has been associated with early gene expression during infection: “[AT][AT][AT][TA]AAAAATGAT[ATG][AG][AC]AAA[AT]”. This motif lacks any of the methylation motifs defined in this study, which may explain the decrease in methylation frequency in the promoter region. However, we found no connection between methylation in the promoter region and temporal gene expression (**Figure S4**). In fact, several genes that are expressed early in the infection cycle, including MCP (**Figure 2A**), have particularly high methylation within the promoter region.

### Nucleotide Bias Surrounding Adenines Influences Methylation

6mA methylation was found to be highly dependent on the nucleotides surrounding the adenine, indicating motif dependent DNA binding typical of MTases (**Figure S5**). Sequence specific analysis revealed that a disproportionate number of methylated adenines occur in the CTAG motif, a 5mer which boasts an average methylation frequency ranging from 34.48 to 57.49%, a stark difference from the average methylation rate of adenines throughout the genome. While methylation frequency at these sites is relatively high, this motif is not common in the viral genome, with the “CTAGG” motif occurring approximately once every 4500 bases, while the analogous “TCAGG” motif occurs once every 360 bases. Likewise, polyadenine regions were largely unmethylated, a pattern that is particularly exacerbated by the presence of one or more adenines downstream of the adenine in question. Classical dam motifs GATC and CATG, which have been identified in the Chlorovirus PBCV-1 and are typically associated with RM systems, bear slight methylation signals at frequencies of 8.3-18.5% and 12.3-14.7% respectively.

To test for co-occurrence of adenine methylation, the methylation frequencies of adenines within ten nucleotides of highly methylated adenines (methylation frequency > 80%) were collectively examined. Sites that are located 2-3 nucleotides upstream and 2 nucleotides downstream on the same strand of the highly methylated adenine are frequently hypomethylated (**Figure S6**). Interestingly, sites that are three and five nucleotides upstream of the highly methylated adenine display elevated methylation frequencies on the opposite strand (**Figure S6**). Biases that exist in favor of nucleotide occurrence surrounding a methylated adenine can also be identified using this approach. When considering only sites surrounding a methylated adenine, the proportion of adenines at 5 and 4 bases upstream and 1 base downstream, and the proportion of guanines 1 base upstream, steadily decreased as a function of minimum methylation frequency (**Figure S7**). Likewise, the proportion of thymine at 5 bases upstream, guanine at 4 bases upstream, and cytosine 1 base upstream, increase (**Figure S7**).

### Several Complex Motifs Identified Display Variable Methylation Levels

To search for more complicated motifs, Nanomotif was used to identify motifs in the genome that were enriched for methylation, using a stringent cut-off of 70% methylation frequency to define methylated sites. Nine motifs were identified between the three sequencing libraries (**Table 1**), including the CTAG motif previously identified. Other derivative motifs to CTAG were also identified, either containing additional nucleotides within (CTNNAG) or flanking the motif (CTAGY). These variations of the CTAG motif represented seven out of nine of those identified, not including GTNNAC and TGNNCA. However, these two motifs, along with CTNNAG, were identified as targeted for methylation in each library.

**Table 1.**
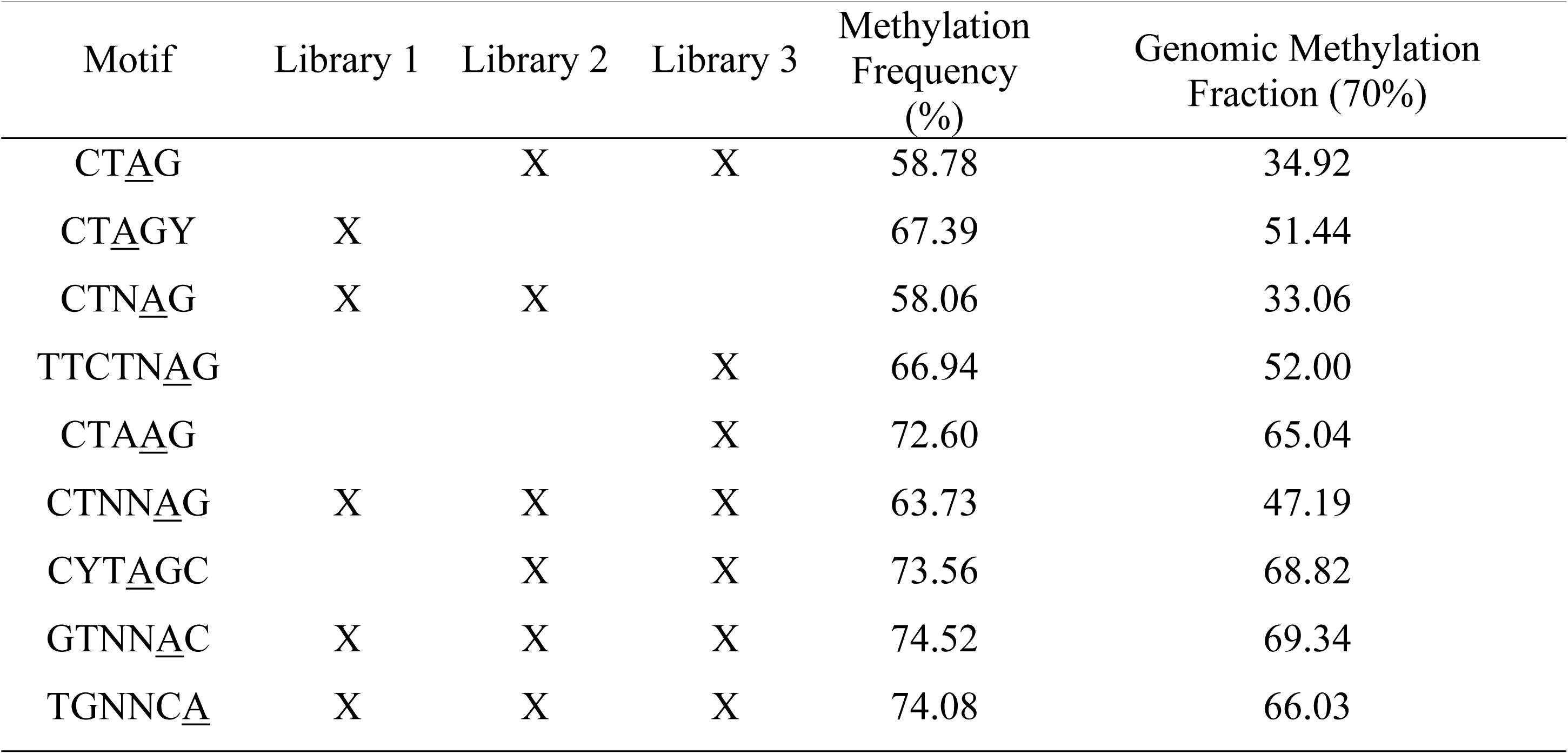
Motifs detected in at least one of three sequencing libraries of AaV. Detection in a library is represented with an X. Methylation frequency and genomic methylation fraction are averaged across all three libraries.

Across the nine motifs, GTNNAC and TGNNCA were also the most highly methylated, with an average methylation frequency of 74.52% and 74.08% respectively. Meanwhile, CTAG and CTNAG exhibited the lowest average methylation frequency of identified motifs, at 58.78% and 58.06% respectively. Collectively, the identified motifs represent a vast majority of the methylated sites, including approximately 74% of sites with a methylation frequency > 50% and approximately 92% of sites with a methylation frequency greater than 70% (**Figure 3A-E)**. Motifs distribution is also largely consistent with highly methylated sites found globally throughout the genome. To rule out false signals that might be attributed to sequencing errors, a WGA library was used to correct methylation frequency scores. Most sites attributed to motifs remained significantly methylated, as compared to classical methylation motifs CATG and GATC (**Figure 3F-G**). Interestingly, while the motifs enjoy a more cosmopolitan distribution, the CATG motif is clearly more abundant at one end of the linearized genome (**Figure 3F**).

**Figure 3.**
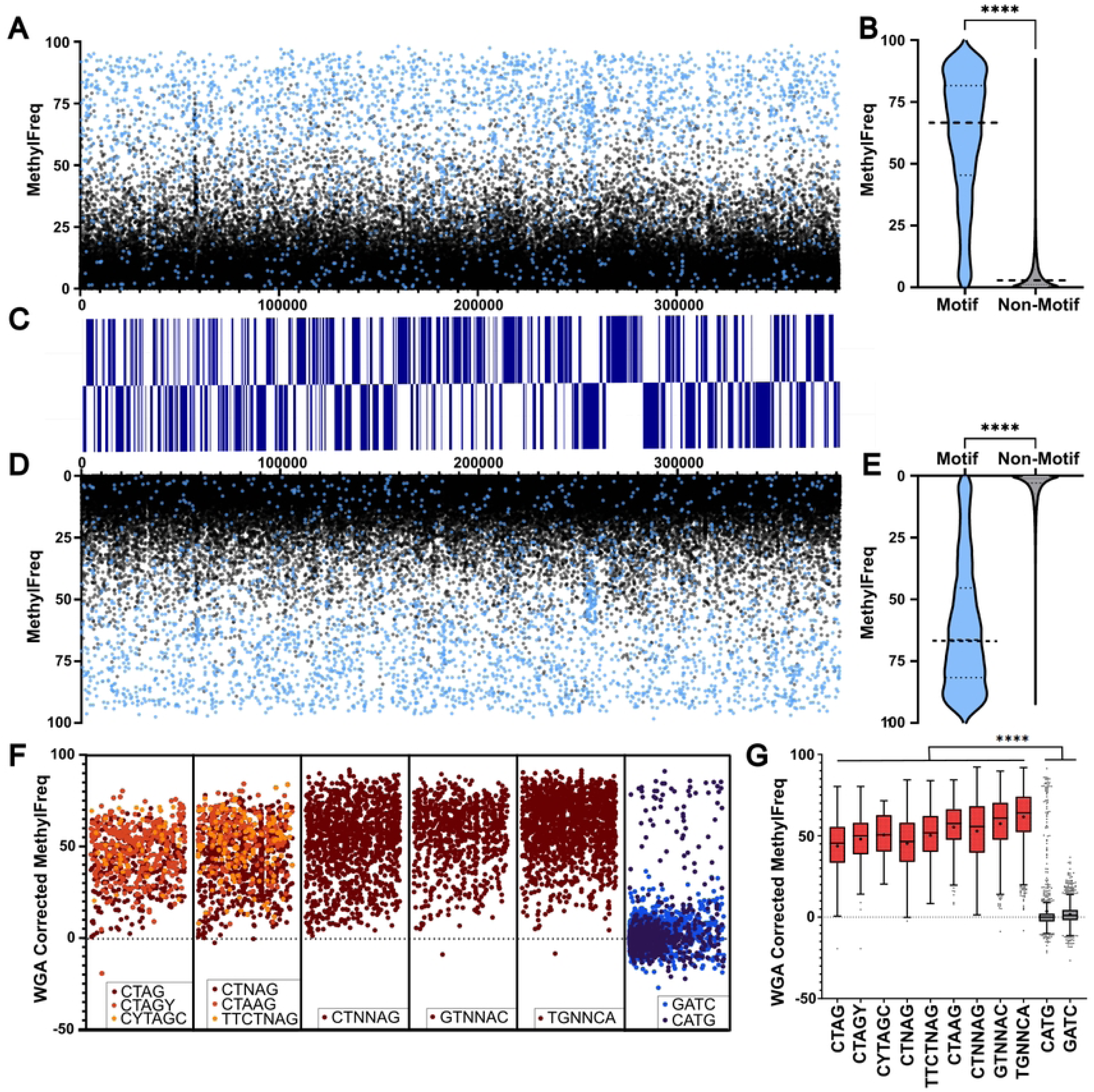
Methylation frequency of characterized 6mA motifs on a genomic scale. Genomic maps show mean methylation frequency for each site on the positive strand (A) and negative strand (D) colored based on whether the site is associated with one of the 9 identified motifs (blue) or not (black). Respective coding regions for each strand are shown in blue (C). Violin plots for each strand detail the differences between motif-associated and non-motif sites on positive (B) and negative (E) strands. Scores for the identified motifs as well as the two classical recognition motifs GATC and CATG were corrected with WGA library scores (F). Each box in F represents a linearized genome meaning each site corresponds to its respective position. Tukey’s box plots were generated to show the distribution of WGA-corrected methylation frequencies, with means denoted as plus symbols (G). ****: p < 0.0001.

Most sites displayed similar distributions regarding variance, with most standard deviations below 10% and barely any above 20% (**Figure 4**). Distribution of methylation frequency means for each individual site varied more depending on the motif, however (**Figure 4**). In certain cases, motifs targeted for methylation contain ambiguous nucleotides (*e.g.* the ambiguous CTNAG) in one library while being defined in other libraries (*e.g.* the specified CTAAG). Comparing the shifts in mean methylation frequency and standard deviation between ambiguous motifs and their respective specified motifs reveals that the specified motifs are both more consistently and frequently methylated (**Figure 4A-B**). Still, a proportion of the highly methylated sites belong to the ambiguous motif only, implying that the specified motif does not completely account for all the associated methylation.

**Figure 4.**
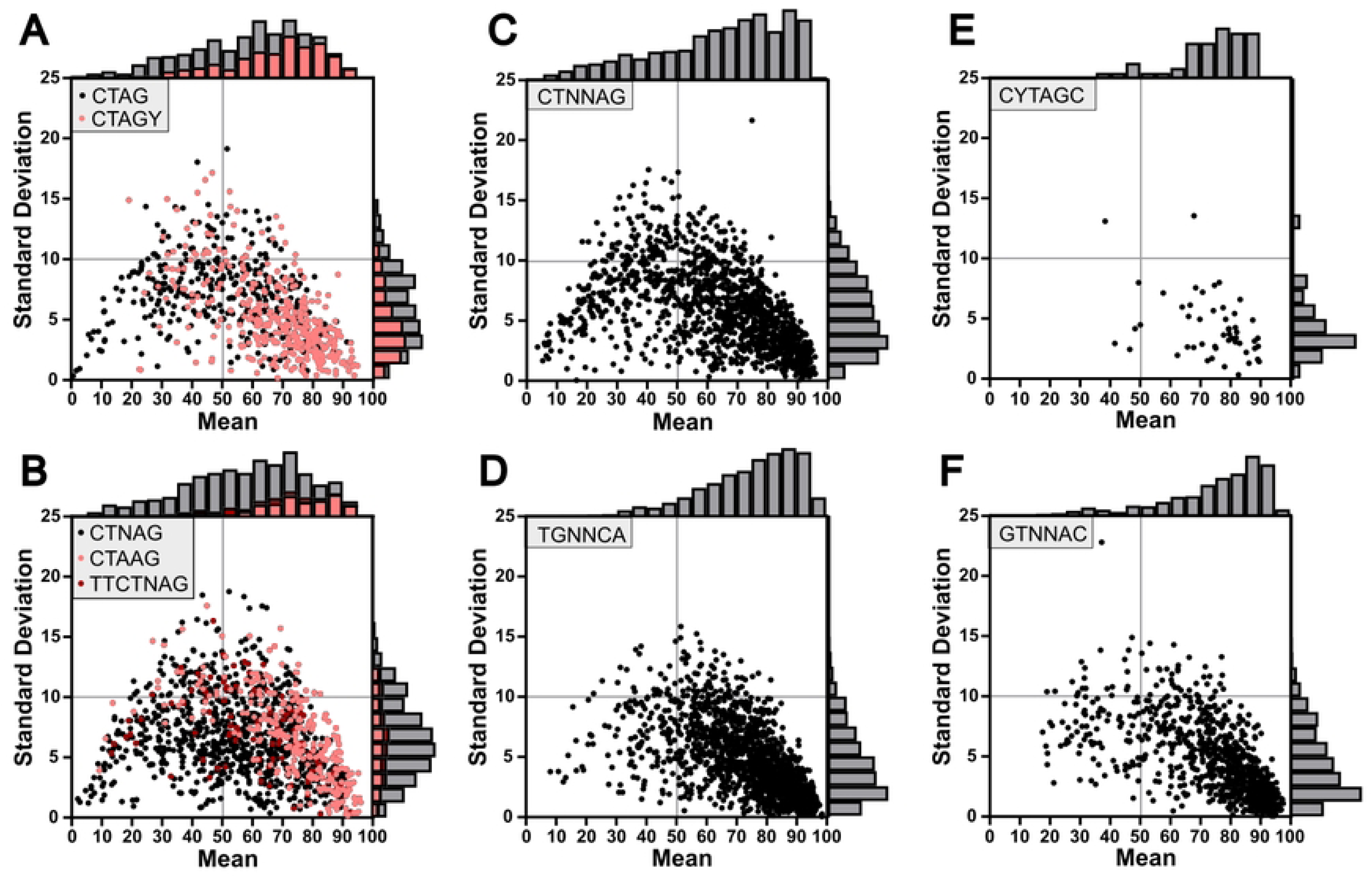
Distribution of methylation frequency for AaV methyltransferase targeted motifs. Distribution of methylation score averages versus standard deviations for each site belonging to a respective motif, with derivative motifs clustered into single plots (A-B). Histograms detail total counts of sites within respective regions.

Considering that five motifs are palindromic (*i.e.* the reverse complement sequence is identical), we sought to determine rates of full and hemi-methylation. To be considered a fully methylated motif, both respective adenines on the positive and negative strand had to reach a minimum methylation frequency of 70%, whereas only one strand being methylated at this level was considered hemi-methylation. The palindromic motifs that are derivatives of the CTAG motif, thus CTAG itself, CTNAG, and CTNNAG, all had low levels of full methylation (<20%) and were hemi-methylated over 50% of the time (**Figure 5A-C**). The GTNNAC and TGNNCA motifs, while still displaying high rates of hemi-methylation, were much more likely to be fully methylated (40-60% of sites) while also being much less likely to be completely unmethylated than the CTAG derivatives (**Figure 5D-E**). In these respective motifs, a significant increase in proportional full methylation was detected as compared to the CTAG derivatives (**Figure 5F-G**).

**Figure 5.**
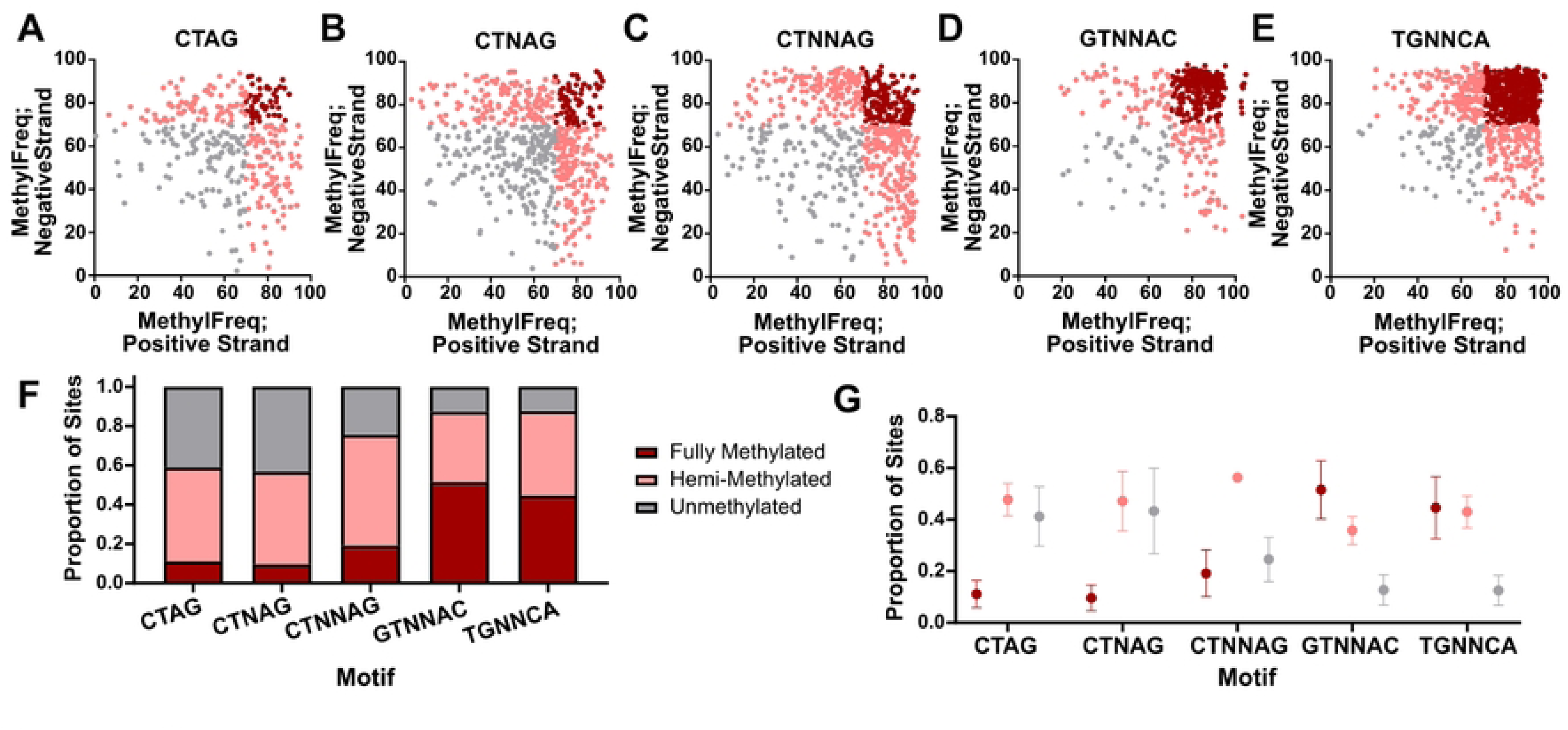
Patterns in reciprocal methylation among palindromic motifs. Reciprocal methylation plots for all palindromic motifs with fully methylated sites displayed in dark red, hemi-methylated sites displayed in pink, and unmethylated sites displayed in gray. Proportion of all sites for each motif (F) and average and the standard deviation of each proportion (G) are denoted.

To further verify the authenticity of the defined methylation motifs, a series of restriction enzyme digests were performed using the enzymes Hpy166II (targeting GTNNAC), XhoI (targeting CTCGAG), and XbaI (targeting TCTAGA). DpnI and DpnII only cleave GATC in which the adenine is either methylated or unmethylated, respectively. Considering GATC sites are apparently unmethylated in AaV, these enzymes were used as negative and positive controls for the restriction digests. None of these enzymes were capable of degrading raw viral DNA, though degradation by DpnII showed the viral genome is definitively not methylated at GATC sites (**Figure S8**). WGA viral DNA was clearly digested by Hpy166II, with potential digestion by XhoI and XbaI as well, compared to the negative control (**Figure S8**). What’s more, *A. anophagefferens* DNA was heavily digested by both Hpy166II and XhoI, implicating divergent methylation patterns between host and virus. As neither XhoI nor XbaI showed strong digestion on AaV or amplified DNA, likely due to the motifs’ infrequency in the viral genome, both were used concurrently. While the two enzymes together showed digestion of amplified DNA, unamplified DNA was unaffected (**Figure S9**).

### Phylogenetics of AaV Methyltransferases May Signify Unique Origins of Methylation Targets

The six DNA MTases encoded by AaV were placed into a phylogenetic tree of *Nucleocytoviricota* MTases which have either been shown to methylate a specific site (17) or are predicted to do so based on homology to defined sequences in REBASE (**Figure 6**). The resulting phylogeny can be broken into four distinct clades, of which three contain AaV MTases.

**Figure 6.**
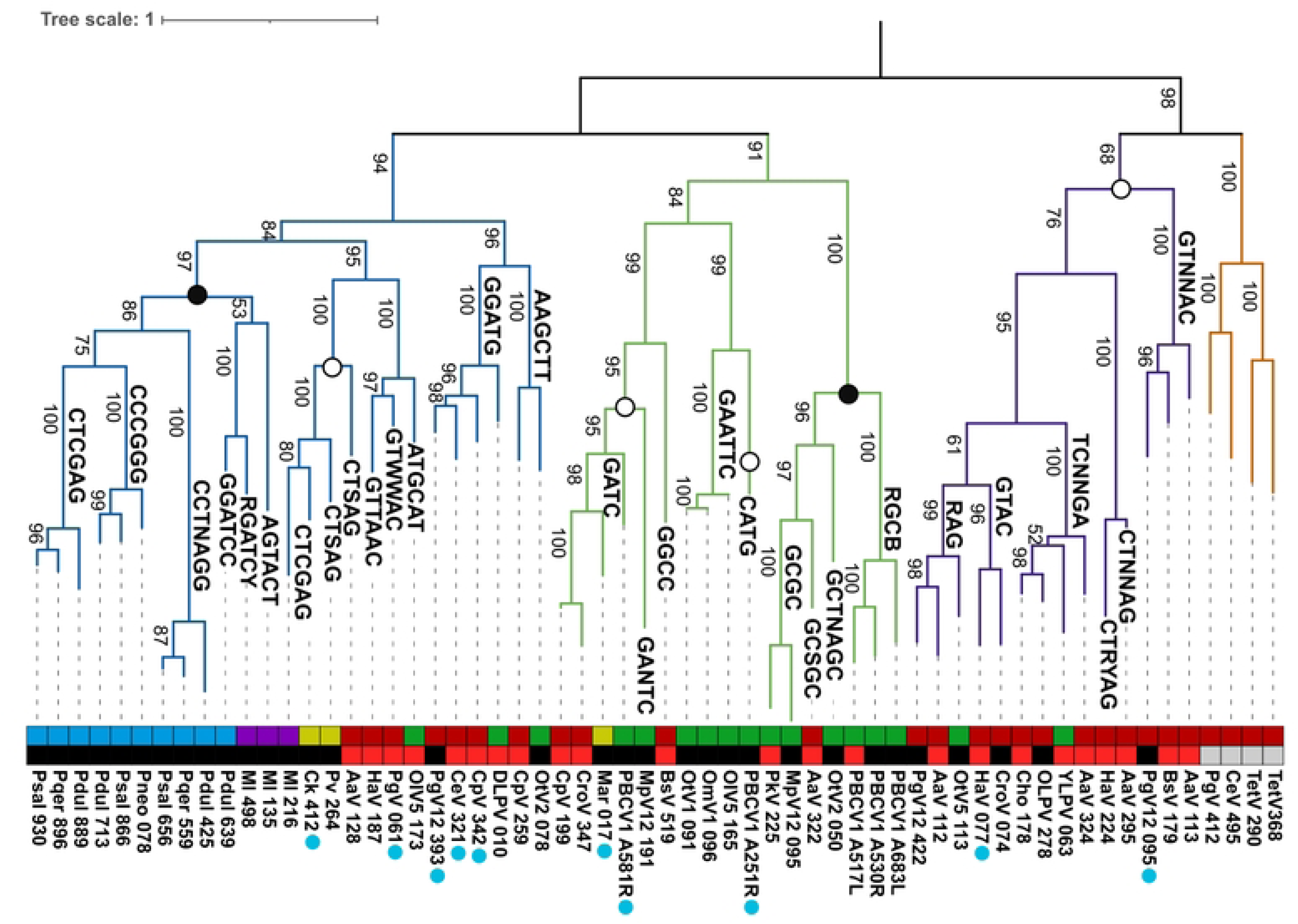
Protein tree of *Nucleocytoviricota* type II DNA methyltransferases. Branches which have a defined or predicted motif are labeled as such. The upper colored bar defines the respective family for each virus while the lower colored bar defines the quality of the predicted target sequence, black meaning the MTase is predicted to target the corresponding sequence by REBASE and red meaning the MTase is predicted to target the corresponding sequence based on homology to other MTases alone. Blue dots below sequence IDs represent the presence of a neighboring restriction endonuclease. Dots on nodes signify the targeted base of all known MTases within the clade, either cytosine (black) or adenine (white). Bootstraps are displayed on branches. Psal: Pandoravirus salinus; Pdul: Pandoravirus dulcis; Pqer: Pandoravirus quercus; Pneo: Pandoravirus neocaledonia; Ml: Mollivirus; Ck: Cedratvirus kamtchatka; Pv: Pithovirus sibericum; AaV: Aureococcus anophagefferens Virus; PgV: Phaeocystis globosa Virus; CeV: Chrysochromulina ericina Virus; CpV: Chrysochromulina parva Virus; DLPV: Dishui Lake Phycodnavirus; EhV: Emiliania huxleyi Virus; MpV: Micromonas pusila Virus; Mar; Marseillevirus; CroV: Cafeteria roenbergensis Virus; OlV: Ostreococcus lucimarinus Virus; OtV: Ostreococcus tauri Virus; OmV: Ostreococcus mediterraneus Virus; HaV: Heterosignma akashiwo Vius; TetV: Tetraselmis Virus; BsV: Bodo saltans Virus.

Clades I and II contain one AaV MTase each (**Figure S10**). Clade I contains one AaV MTase (AaV_128) which is predicted to target the CTSAG motif based on REBASE homology. This is also the only type II β MTase encoded by AaV, as all others are of the γ subclass. Despite other *Imitervirales* and *Algavirales* methyltranferases in this clade, the AaV MTase bears relatively increased similarity (approximately 35%-40% amino acid similarity) to those of Cedratvirus, Pithovirus and *Mollivirus sibericum*, which again target either a CTNNAG motif derivative or CTNAG motif derivative. Notably, while REBASE initially predicted that the Cedratvirus MTase would target CTSAG as well, PacBio sequencing revealed that it instead targets the CTCGAG motif (17). This may imply that the related AaV MTase either targets the CTNNAG motif or the CTNAG motif.

Interestingly, Clade I contains a cluster of cytosine specific MTases (subclass IA; **Figure S10**), which belong to either the Pandoraviruses or Molliviruses. A monophyletic sub-clade of these genes belongs exclusively to the Pandoraviruses with zero homologs outside of this specific viral family. While these enzymes perform 5mC methylation (17), two of the described motifs mirror the CTNAG and CTNNAG motifs identified in the AaV methylome. In fact, the CTCGAG motif appears in both several Pandoraviruses as well as *Cedratvirus kamchatka*, though cytosine and adenine are targeted respectively between the two viral MTases.

Clade II is primarily comprised of MTases of the *Algavirales*, among which is the PBCV-1 gene M.CviAII, which targets the adenine in the CATG motif and M.CviAII which targets the adenine in the GATC motif (38). Despite this, this clade varies in both the predicted motif for methylation as well as type of methylation (6mA or 5mC). Still, distinct lineages including the Marseilleviruses and the *Imitervirales* are present throughout this clade. The truncated cytosine-specific MTase AaV_322 falls within subclade IIC (**Figure 6, Figure S10**), which has 100% bootstrap support as distinct from the adenine-specific enzymes in this clade.

Clade III contains the four remaining AaV MTases, though they cluster into three distinct subclades supported by high bootstrap values (**Figure 6, Figure S10**). Notably, all defined MTases in this clade are predicted to target adenine. Two AaV MTases are in subclade IIIA, which are predicted to target an ambiguous RAG motif and a TCNNGA motif. The MTases identified in this clade belong to both *Algavirales* and *Imitervirales* and do not group taxonomically. While the RAG targeting MTase may be unclear, the TCNNGA targeting MTase may be instead represent the methylation of the TGNNCA motif. The only associated MTase that is defined by REBASE is that of the Organic Lake phycodnavirus, which is only characterized based on the prototype restriction endonuclease Hpy178III.

Subclade IIIB and IIIC contain the final two AaV MTases which are predicted to target the GTNNAC motif and the CTNNAG motif. The MTases identified in these subclades exclusively belong to the *Imitervirales* specifically belonging to the Mesomimiviridiae, including Heterosigma akashiwo Virus, Phaeocystis globosa Virus, and Bodo saltans Virus. While the defined motifs for these viruses are largely based on homology alone, the further appearance of the experimentally verified motifs from the AaV methylome in these clades is a strong indication of their targeted sequence. With this information in mind, the expression level of AaV MTases during infection was identified from a previous study (39). Notably, the cytosine specific MTase was expressed at the highest level within 12 hours of infection (**Figure S11**). Of the adenine-specific MTases, the gene predicted to target the GTNNAC region was also highly expressed at the 12 hpi and the highest expressed MTase at 23 hpi (**Figure S11**).

### Genes Cluster Based on Enrichment of Specific Motifs

To determine whether the identified motifs were enriched in ORFs encoded by AaV, we used the DistAMo web tool to map the frequency of each motif relative to genomic position. Genes with a z-score greater than two were identified for each motif and annotated functionally from the AaV reference genome (**Table S3**). Genes were then clustered in Cytoscape with a single connection between a gene and a motif signifying overrepresentation of the given motif in said gene (**Figure 7**).

**Figure 7.**
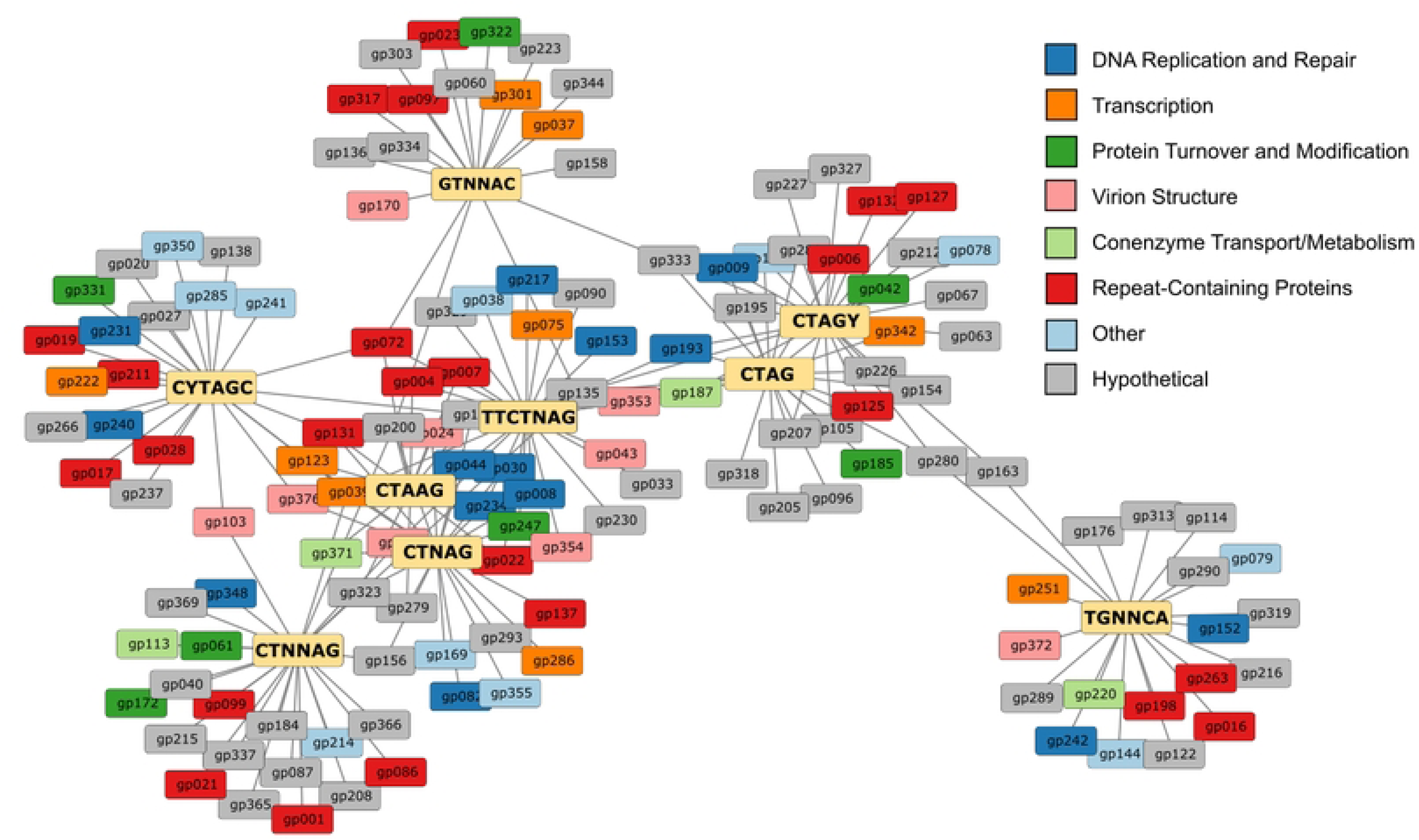
Gene clusters formed based on linkages between motif enrichment. Characterization for motif enrichment within a coding region was determined based on DistAMo results. Only genes with a z-score for motif enrichment greater than 2 were considered. Each motif (yellow) is connected to any genes that are enriched with the given motif. Different COG classifications of genes are denoted by color.

Overall, 135 genes (35% of all predicted coding regions in AaV) were enriched with at least one of the identified motifs. Each individual motif is enriched within 18 to 25 genes. 55 of the overrepresented genes (41%) could be functionally annotated, which is elevated from the full genome of AaV in which only ∼25% of genes can be assigned to functional clusters of orthologous groups (COGs). While there was no obvious relationship in function for genes clustered around a specific motif, several important COG categories were represented across all motifs. Fourteen genes related to DNA replication and repair were identified, including several DNA polymerases, along with nine related to transcription and another nine related to virion structure, including the major capsid protein (**Table S3**).

While certain genes are often enriched with multiple motifs, most clustering appears to primarily occur between closely related motifs (*i.e.* CTAG and CTAGY). Motifs like CTNNAG, CYTAGC, GTNNAC, and TGNNCA are largely isolated with few connections to other clusters. Repeat-rich proteins are also detected in every cluster, many of which belong to the domain of unknown function (DUF) 285 family. While these proteins are present in the same family, their genetic makeup seems to differ heavily, with one motif repeating dozens of times and none of the other motifs being detectable.

## DISCUSSION

Laboratory studies as well as metagenomic and metatranscriptomic data from natural systems have yielded remarkable discoveries among the “giant viruses” (40, 41). Yet beyond nucleotide sequence, little is known about their genomes with respect to the relevance of epigenetic modification of *Nucleocytoviricota* DNA. This is of particular importance considering the wide range of *Nucleocytoviricota* species as well as the high density of DNA MTases encoded by certain viruses relative to the total genomic coding potential (see **Figure 1B**). Methylation of viral DNA has been shown to influence the number of infectious viral particles produced during lysis (42) and allows for resistance to DNA digestion via endonuclease activity (16, 17). Given DNA methylation has also been ascribed to virus-resistance mechanisms in both bacteria (11) and eukaryotes (43), the physical proximity between viral and host genetic material may serve as a means for acquisition and repurposing of host MTases to allow for invading genome survival. Thus, characterization of methylation profiles across viral genomes may improve our ability to predict infection outcomes.

Annotation of a given sequence’s methylation state is generally untethered from traditional sequencing approaches: it has historically required lengthy, multi-step analyses like whole genome bisulfite sequencing and restriction enzyme-based analyses. Yet advancements in long-read sequencing have provided the ability to detect methylation through traditional library preparation techniques (44). Moreover, analysis with the Nanopore MinION Flongle attachment allows for fiscally reasonable high throughput genotyping of small bacterial and viral genomes with both high genomic coverage and read length (45). This can then allow for the assembly and subsequent read mapping to a complete genome, providing high-confidence methylation scores for adenines and cytosines in a genome (46).

To further our understanding of how giant viruses utilize MTases, we completed a comprehensive genomic sequencing approach of the algal virus *Kratosvirus quantuckense* strain AaV using Nanopore long-read sequencing. By sequencing the AaV genome in biological triplicate, we have provided a perspective on not only the methylation state of the viral genome, but also the consistency of methylation across multiple generations of packaged viral DNA. From this we defined nucleotide biases that influence the targeted sequences for methylation and characterize several unique DNA methylation motifs which can be loosely ascribed to specific AaV MTases. Based on the distribution of methylation across the AaV genome, we suggest genomic methylation may serve as both functional and ancestral traits in *Nucleocytoviricota* genomes.

### Detection of AaV Methylation is Highly Repeatable

Adenine and cytosine methylation of the AaV genome was consistent, with standard deviations for replicate infections of host cultures consistently below 10% for most sites. Considering different environmental conditions can alter genomic methylation (2, 47), the homogeneity of the highly methylated sites serves as an important baseline for future studies. The relationship between mean methylation frequency and standard deviation for a given site prefers to follow a quadratic association, with the highest amount of variation detected in the sites that average a moderate methylation frequency, around 50%. This suggests that the sites responsible for the highest methylation frequency scores are universally targeted for methylation, at least under the conditions tested here. From a consistency standpoint, this may indicate a strong selective pressure for methylation of these sites for the propagation of viral progeny. Meanwhile, the sites with moderate levels of methylation may be inconsistently methylated due to a lack of the same pressure, possibly the result of non-specific binding of an MTase to a site other than its preferred recognition domain.

While most nucleotides are definitively unmethylated across the AaV genome, adenine-specific methylation clearly displays higher methylation frequency and genomic methylation fraction compared to all cytosine-specific methylation. This may be attributed to the sheer number of MTases predicted to be adenine-specific (five) compared to only a single cytosine-specific MTase. However, the fact that, within 12 hours of infection, the normalized read abundance of this cytosine-specific MTase is often more than double that of the adenine-specific MTases (**Figure S11**), reveals a false equivalence. Higher MTase expression does not correspond to increased methylation levels. Thus, it is likely that the cytosine-specific MTase is either extremely inefficient or provides an additional function for the virus during infection, possibly in nucleotide metabolism or even methylation of the host cytosines as a potential gene silencer (48). We note that the host *A. anophagefferens* also encodes cytosine-specific MTases, and thus methylated cytosine residues on the viral genome may be a result of host MTase activity.

Spatial distributions of highly methylated sites imply a functional role in this activity among both cytosines and adenines. Several genome regions are heavily enriched in highly methylated cytosines, with these areas primarily being repeat-dense. This may be representative of altering the steric hindrance around these sites to influence the formation of tertiary structures in the DNA strand during synthesis or affect the binding of transcriptional proteins (49, 50). Likewise, the significant decrease in methylated adenines in the region immediately upstream of coding regions may imply that methylation of viral DNA can act as a deterrent to transcription initiation, at the very least. Methylation shifts in the promoter region are highly common in many eukaryotic systems (1, 4). However, as there are no linkages between promoter region methylation frequency and time of expression during infection, it does not restrict transcription. For example, the major capsid protein, which is expressed within the first five minutes of infection, contains an adenine methylated at approximately 70% immediately upstream of the start codon.

### Targeted Methylation is Influenced by Divergent Nucleotide Motifs

While initial attempts at identifying nucleotide biases surrounding methylated sites revealed several preferred nucleotide pentamers, particularly in association with the “CTAGN” motif, very few pentamers had truly high methylation frequencies. Further motif analysis revealed that this is likely the effect of many of the defined motifs containing multiple ambiguous nucleotides. Among the motifs identified, GTNNAC and TGNNCA were targeted at the highest methylation frequency and genomic methylation fraction, two motifs that have largely not been identified as targets for viral DNA methylation. CATG and GATC, two motifs functionally characterized as methylation targets in PBCV-1 (16), were unmethylated, for the most part, pointing to a unique ancestry of MTases in AaV as well as the possibility of a diversified function.

Seven motifs identified within the AaV methylome ascribe to a CTAG-like structure, with generic and specific forms being identified across the three sequencing libraries. These include the generic CTAG being specified into CTAGY and CYTAGC and the generic CTNAG being specified into CTAAG and TTCTNAG. In all cases, specification of a motif yielded an increase in average methylation frequency and genomic methylation fraction, though specification did not account for all the highly methylated sites of the original generic motif. This may be an indicator of a “leaky methylation” phenotype. As such, there are likely five adenine-specific DNA MTases encoded by the AaV genome, despite the presence of nine identified motifs. This suggests that some MTases are responsible for methylation of multiple motifs. Thus, a MTase may have an optimal DNA recognition sequence which ends up very consistently methylated (*i.e.,* CTAAG), but may also non-specifically bind to other similar derivatives of the same motif, explaining why CTNAG is frequently recognized by motif detection software. Given seven of the nine identified motifs possess at least one ambiguous nucleotide, it is possible that many of the AaV MTases have become leaky with time, allowing for more varied methylation across the genome.

Nucleotide proportion as a function of minimum methylation frequency displays unique distribution depending on the nucleotide and site in question, in some cases appearing largely linear while in other cases displaying polynomial distributions (**Figure S7**). This could be reflective of the motif that is causing the distribution. For instance, none of the identified motifs contain adenine immediately following the 6mA site, explaining the rapid decrease in adenine proportion at that site. Meanwhile, a thymine five bases upstream of the 6mA is likely to belong to the TGNNCA motif, which could also be said regarding a cytosine one base upstream. Likewise, a guanine four bases upstream of the 6mA likely represents the GTNNAC motif.

### Phylogenetics and Genomic Distribution of Motifs Reveals Importance of Methylation in AaV

Phylogenetic analysis of the AaV adenine-specific DNA MTases has allowed for the association with several possible DNA recognition sequences which correspond to previously identified motifs. The clade I MTase likely targets CTNAG or CTNNAG, the clade IIIC MTase likely targets GTNNAC, and the clade IIIB MTase likely targets CTNNAG. While one clade IIIA MTase is predicted to target TCNNGA, this is instead most likely representative of the TGNNCA motif which was identified in the AaV methylome. The other clade IIIA MTase is predicted to target an RAG sequence, which is unlikely to be an actual motif targeted by any AaV MTases due to its simplicity and abundance in the AaV genome. It is unclear whether this MTase is representative of one of the identified motifs, as many of them contain an adenine followed by a guanine, or whether it instead targets a motif that was unidentified by our analysis.

The presence of a MTase in clade I is particularly intriguing considering there are seemingly no closely related *Mimiviridae* MTases and this gene is more phylogenetically like those of the Pithoviruses and Molliviruses than anything else. The origin of such an MTase is thus perplexing, though these may have been acquired from an ancestral eukaryotic host.

Additionally, confounding is the lack of type II MTases encoded by the classical *Mimiviridae* including Acanthamoeba polyphaga Mimivirus (APMV), Moumouvirus, and Megaviruses (51, 52). While the closely related “extended *Mimiviridae”* or *Mesomimiviridae* including AaV appear to often encode multiple DNA MTases, these amoebal viruses are apparently unmethylated (17). This seems to suggest that these genes can be lost, at least in the context of some host backgrounds, though most algal viruses appear to encode at least one MTase.

Though adenine-specific motifs are generally distributed evenly throughout the AaV genome, they are overrepresented in certain areas. Identifying genes located in these areas reveals that certain genes may be more likely to be affected by methylation of some motifs as compared to others. Clustering of genes based on their connection to the nine defined motifs showed that while there is an overlap between closely related motifs, divergent motifs largely affect different genes.

Individual clusters of genes were not found to share any connections based on gene ontology or ancestral source (*i.e.* viral or host-derived). However, the formation of clusters does suggest that in many cases there is not a functional redundancy introduced by the five AaV adenine MTases. In accordance with the phylogenetic analysis, it is unlikely that the MTases target the same motif and thus they end up targeting different regions of the genome as well as affecting different genes. The heterogeneity of MTases in this virus mirrors that of certain bacteria, including *Helicobacter pylori* and *Microcystis aeruginosa*, which may encode between 20 and 50 methylation systems in a single genome (11, 53, 54). This creates a paradigm in which the expression of all MTases during infection is important, as absence of one of the MTases may leave certain areas undermethylated. Whether this would result in improper packaging of viral DNA, degradation by host endonucleases, or shifted gene expression is yet unknown, but the prevalence and distribution of methylation across the viral genome still implies functional significance.

### Concluding Remarks

Beyond defining the methylome of AaV, our analysis has shown that the annotation and characterization of methylation on a large viral genome can be performed at a relatively low cost and high efficiency. We sequenced the AaV genome a total of four times in this analysis, which yielded three characterized methylomes which displayed high consistency. Our characterization was further justified when WGA-sequencing showed that amplification of the AaV genome (i.e., a negative control) quenched the methylation signal reported in the pipeline, supporting our consensus that the sites described are in fact methylated and are not a byproduct of the sequencing technology used. Collectively this base methylome for the virus provides the opportunity for research into the factors that influence methylation and the downstream effects of differential methylation. Nutrient stress has been associated with changes in methylation in some plant species, and reducing the activity of MTases during infection may reveal more about the nature of methylation in *Nucleocytoviricota* infection. Similar observations like those made in plants as well as other mechanistically focused studies moving forward can now consider how genomic modifications can constrain biological function and/or success beyond the base-code of a genome.

## Acknowledgements

We thank Brittany Zepernick, Robbie Martin, Tim Sparer, Brad Binder, Todd Reynolds, Katelyn Houghton, Laura Smith, and Kennedi Hambrick for discussions regarding this work.

## Data and Materials Availability

All sequencing data generated in this study have been submitted to NCBI under the BioProject accession PRJNA1292256 as sequence read archives with the accession numbers SRR34564230-SRR34564239. All python scripts used in this study are available at https://github.com/Wilhelmlab/Truchon2025.

